# Synthetic and biological surfactant effects on freshwater biofilm community composition and metabolic activity

**DOI:** 10.1101/2021.07.30.454494

**Authors:** Stephanie P. Gill, William R. Hunter, Laura E. Coulson, Ibrahim M. Banat, Jakob Schelker

## Abstract

Surfactants are used to control microbial biofilms in industrial and medical settings. Their known toxicity on aquatic biota and their longevity in the environment has encouraged research on biodegradable alternatives such as rhamnolipids. While previous research has investigated the effects of biological surfactants on single species biofilms, there remains a lack of information regarding the effects of synthetic and biological surfactants in freshwater ecosystems. We conducted a mesocosm experiment to test how the surfactant sodium dodecyl sulphate (SDS) and the biological surfactant rhamnolipid altered community composition and metabolic activity of freshwater biofilms. Biofilms were cultured in the flumes using lake water from Lake Lunz in Austria, under high (300 ppm) and low (150 ppm) concentrations of either surfactant over a four-week period. Our results show that both surfactants significantly affected microbial diversity. Up to 36% of microbial operational taxonomic units were lost after surfactant exposure. Rhamnolipid exposure also increased the production of the extracellular enzymes, leucine aminopeptidase and glucosidase, while SDS exposure reduced leucine aminopeptidase and glucosidase. This study demonstrates that exposure of freshwater biofilms to chemical and biological surfactants caused a reduction of microbial diversity and changes in biofilm metabolism, exemplified by shifts in extracellular enzyme activities.

**Key Points:**

1. Microbial biofilm diversity decreased significantly after surfactant exposure.
2. Exposure to either surfactant altered extracellular enzyme activity.
3. Overall metabolic activity was not altered, suggesting functional redundancy.

## Introduction

In aquatic ecosystems, microorganisms live preferentially in microbial biofilms and act as major drivers of the biogeochemical cycling of carbon, nitrogen, and other nutrients (Paul *et al*. 1991; Decho 2000; Dang and Lovell 2016; Flemming *et al*. 2016; Battin *et al*. 2016). In streams and rivers, microbial biofilms play an important role in the carbon cycle (Lyon and Ziegler 2009; Qu *et al*. 2017). Microbial biofilms resident in fluvial sediments can also substantially reduce inorganic nitrogen (N) loads in N-rich aquatic ecosystems through denitrification (Zhang *et al*. 2016; Cui *et al*. 2017). In N-limited aquatic systems, they may help to facilitate the transfer of N to higher trophic levels (Dodds *et al*. 2000).

While microbial biofilms play an important ecological role, they are of concern in industrial and medical applications. For example, biofilms can facilitate the growth of antibiotic resistant bacteria (Lindsay and von Holy 2006; Srey *et al*. 2013; Bowler *et al*. 2020). Synthetic surfactants, such as sodium dodecyl sulphate (SDS), are widely used in personal care and cleaning products. They reduce biofilm growth in a wide range of industrial, medical, and environmental contexts (Waaler *et al*. 1993; Landa *et al*. 1999; Rigotti *et al*. 2017; Sloup *et al*. 2016; Ueda *et al*. 2019). Surfactants are amphiphilic chemicals that reduce surface tension within a liquid, sometimes through the formation of micelles. Micelles occur when molecules form an aggregate with hydrophobic tails on the inside (Schmitz 2018) and hydrophilic heads on the outside of an amphiphilic compound (Milovanovic *et al*. 2017).

Research on the effects of surfactants on microbial biofilms has primarily focused on single-species biofilms or those commonly found in medical and industrial environments. Direct applications of the anionic synthetic surfactant SDS has shown significant disruption in the growth and development of *Pseudomonas aeruginosa* biofilms (Díaz De Rienzo *et al*. 2016; Nguyen *et al*. 2020). Effects on other single species or medical biofilms have also been investigated including methicillin-resistant *Staphylococcus aureus* (Ueda *et al*. 2019), *Pseudomonas fluorescens* (Simoes *et al*. 2008), and *Salmonella enterica* serovar *typhimurium* (Mireles II *et al*. 2001). Inhibited biofilm formation was seen to occur within all investigations.

The use of synthetic surfactants has been questioned due to the potential environmental impact, specifically the longevity and toxicity to aquatic biota. For example, previous research has shown that synthetic surfactants, such as Triton X-100 (Octylphenol Ethoxylate), are not easily biodegradable under aerobic nor anaerobic conditions (Mohan *et al*. 2006). Other synthetic surfactants such as SDS have also been noted as toxic to a variety of microorganisms such as *Vibrio fischeri*, fish and shellfish such as *Clarias batrachus* and *Mytilus galloprovincialis*, yeast, and also small invertebrates, such as *Daphnia magna*, and *Tigriopus fulvus* (Mariani *et al*. 2006; Lima *et al*. 2011; Franzetti *et al*. 2012; Messina *et al*. 2014; Cao *et al*. 2020; Kumar *et al*. 2020). Research has therefore shifted towards identifying a natural alternative that may be less harmful to the aquatic environment.

Biofilm-dwelling microorganisms naturally produce a number of surfactants to control the surrounding extracellular polymeric substance (EPS) matrix. The most common substances of this group are rhamnolipids and sophorolipids which are glycolipids formed by the combination of either a long-chain aliphatic acid or hydroxyaliphatic acid with a carbohydrate (Desai and Banat 1997, Marchant and Banat 2012). When exposing single-cell biofilms of *P. aeruginosa* to rhamnolipids, growth and development were disrupted, and cell viability was reduced (De Rienzo *et al*. 2015). Disruptions by rhamnolipids have also been shown for single species biofilms of *Staphylococcus aureus* (Quinn *et al*. 2013; Aleksic *et al*. 2017; Ceresa *et al*. 2021a; Tambone *et al*. 2021), *Bacillus pumilus* (Dusane *et al*. 2010), *and Rhodococcus erythropolis* (Schreiberova *et al*. 2012). Yet, rhamnolipids are biodegradable in a variety of environments, including under aerobic and anaerobic conditions (Mohan *et al*. 2006; Naughton *et al*. 2019; Ramos da Silva *et al*. 2019). Biosurfactants in general have been reported to act as an antibiofilm agents (Banat *et al*. 2014). Given that previous research primarily addressed the effects of biological surfactants on single species biofilms of bacteria relevant to medical settings, there remains a knowledge gap on how the exposure of multi-species freshwater biofilms to surfactants affects biofilm microbial community composition and function.

There is a growing body of evidence that anthropogenic trace contaminants, and contaminants of emerging concern (CECs), such as pharmaceuticals, herbicides, and antimicrobial agents can have major impacts upon the composition and functioning of aquatic microbial communities (Rosi-Marshall *et al*. 2013; Argudo *et al*. 2020; Harjung *et al*. 2020; McClean and Hunter 2020; Ke *et al*. 2020; Kusi *et al*. 2020). Exposure to contaminants such as the antibacterial agent Triclosan (Hay *et al*. 2001; Drury *et al*. 2013) and the herbicide Diuron (Sumpono *et al*. 2003; Seghers *et al*. 2003) have been observed to place microbial communities under a selective pressure that favors microorganisms that can use these contaminants as an additional carbon source. Given that levels of surfactants and their metabolites are consistently among the top contaminants identified within wastewater influent and groundwater (Lara-Martín *et al*. 2014; Schaider *et al*. 2014), it is reasonable to infer that they could represent a major control upon the microbial communities within human affected inland waters.

This study investigated the effects of the synthetic surfactant SDS and the biological surfactant rhamnolipid on microbial biofilm community composition, microbial diversity, and metabolic activity. Using recirculating flume mesocosms, we aimed to answer the following overarching scientific question: What are the effects of a synthetic and biological surfactant on a natural aquatic microbial biofilm, and how do these effects compare? We hypothesized that biofilm exposure to a surfactant will decrease bacterial biodiversity, as the surfactants impact microbial community composition and causes the loss of sensitive bacterial taxa as previous research has shown that chemical exposure can place microbial communities under selective pressure. Secondly, we hypothesize that the resultant changes in microbial community composition will alter biofilm metabolic activity. We planned to test these hypotheses by quantifying changes in community composition through 16s rRNA analysis and changes in biofilm structure and metabolic activity through different analytical methods, such as changes in gross primary productivity (GPP), community respiration (CR), ash free dry mass (AFDM), Carbon: Nitrogen ratios (C:N), and extracellular enzyme activity (EEA).

## Materials and Methods

### Experimental Set up

We set up 15 recirculating experimental flumes near Lunz Am See, Austria following the method described by Roche *et al*. (2017). Each flume was 2.5m long and had an approximate slope of 0.8%. In each flume, ∼5L of water recirculated, which was renewed with unfiltered lake water twice per week. We added rhamnolipid (RL) (R90-100G, Sigma-Aldrich, Darmstadt, Germany) and SDS (sodium dodecyl sulphate, Sigma-Aldrich, Darmstadt, Germany) as treatments of 150 and 300 ppm to three flumes each (hereafter referred to as either RL 150, RL 300, SDS 150, or SDS 300), while three flumes acted as controls. After the first week, 5 µg L^-1^ of phosphorus in the form of Na_2_HPO_4_ was also added to help growth due to low phosphorus in the utilized lake water. Surfactant concentrations were chosen to represent the effective dose of rhamnolipid, as well as double that concentration. Due to malfunction of one flume, *n* for the rhamnolipid treatments was reduced to 2. Treatments were randomly placed among flumes. Artificial light was spread equally over all flumes and light density averaged 9.5 µmol m^-2^ s^-1^ and was the maximum even light density achievable with the provided light sources. We used a light cycle of 12:12 and 18:6 (Cheah and Chan 2021) hours of light versus dark early and later in the experiment to mimic natural light cycles at first and then to encourage biofilm growth later in the experiment. Each flume was equipped with microscope slides (6) and clay tiles (8) to quantify biofilm growth.

We measured pH, dissolved oxygen (DO) (mg/L), water temperature (°C), and flow (by bucket measurement) every other day to ensure stable conditions. Twice per week we analysed DOC, ammonium (NH_4_), NO_x_, and PO_4_ of flume water to ensure they remained stable. In addition, water temperature was monitored by automatic loggers in drainage tanks. After 32 days the experiment was stopped due to adequate biofilm growth observed, and biofilm samples were retrieved from the flumes for further analysis.

### Sample Collection

Two microscope slides were removed from each flume and placed in 120mL Schott bottles with oxygen sensors. Bottles were filled with lake water run through glass fiber filters to remove any microorganisms that may alter the results of the analyses and sealed to remove all air bubbles. Three control bottles with no slides were also set up to test for potential activity within the filtered lake water with control activity subtracted from treatments during analyses. All bottles were placed in an Imago 500 Controller Environmental Chamber (Snijders Scientific B.V., Tilburg, Netherlands) using a 12:12 light cycle for 72 hours. Oxygen measurements were taken at the start and end of each light and dark cycle to obtain data on GPP and CR.

We determined AFDM by weighing 2 clay tiles per flume after drying (9h at 70°C), and combustion (4h at 450°C). AFDM was obtained by subtracting the remaining ash weight from the dry weight. Two tiles from SDS 150 flumes were damaged during drying, and AFDM could not be determined. Biofilms covering two tiles per flume were then scraped into sterilized and pre-weighed 2mL Eppendorf tubes using a sterile razor blade, lyophilised (-0.001mbar; -76□) (Martin Christ, Gefriertrocknungsanlagen GmbH, Osterode am Harz, Germany) and then analysed on a Thermo Scientific FLASH 2000 Elemental Analyser (Thermo Fisher Scientific GmbH, Bremen, Germany) to quantify carbon (% C) and nitrogen (% N) content. Quantities of C and N by mass were then examined in ratios to identify changes in ecological stoichiometry (the relationship between elements, energy, and organisms and their interactions in an ecological system (Sterner and Elser 2002; Van de Waal *et al*. 2018)).

EEA of β-D-1,4-glucosidase (glucosidase), phosphatase, and leucine-aminopeptidase were measured in the biofilms following the method described in Romaní and Sabater (2001). Four mL of sterile water was added to approximately 0.5 g of biofilm per flume that was scraped from the tiles. The mixture was homogenized by shaking. EEA was measured via artificial substrates: Glucosidase and phosphatase using 4-methylumbelliferyl (MUF)-substrates (MUF-β-D-glucopyranoside and MUF-phosphate, respectively (Sigma-Aldrich, Darmstadt, Germany)) and leucine-aminopeptidase using a 7-amino-4-methylcoumarin (AMC)-substrate (L-leucine-AMC, Sigma-Aldrich, Darmstadt, Germany). Artificial substrates were added to the homogenate at a final concentration of MUF-glucosidase; 135 mg L^-1^, MUF-phosphatase; 102.46 mg L^-1^, and AMC-leucine-aminopeptidase; 129.94 mg L^-1^ and incubated for 1 hour in the dark. Fluorescence was measured at the beginning and end of the incubation with a plate reader (Varioskan Flash, Thermo Fischer Scientific, Vaanta, Finland) at 365/455 nm excitation/emission wavelengths for the MUF-substrates and 380/440 excitation/emission wavelengths for the AMC-substrate.

Two microscope slides per flume were scraped using sterile procedures into a sterile 1.5mL microcentrifuge tube for 16s rRNA analysis. DNA was immediately extracted from collected samples using the Dneasy Power Soil Kit (Qiagen GmbH, Hilden, Germany) following the manufacturer’s methodology. We included internal replicates per flume (*n*=2), and experimental controls were used to control for any potential unaccounted sample contamination. Extracted products were sent to LGC Biosearch Technologies (LGC Genomics GmbH, Berlin, Germany). DNA was quantified using gel electrophoresis before amplifying the V3-V4 region of the 16s rRNA gene with polymerase chain reactions using 341F and 785R primers. Resultant DNA were sequenced using the Illumina Mi-Seq platform (300 bp paired-end reads) for amplicon creation.

We used Mothur version 1.44.3 (Schloss *et al*. 2009), and the Mothur MiSeq SOP (Kozich *et al*. 2013) with some modification, to analyze returned reads. A maximum homopolymer of 8 was chosen due to the limitation of the reference database (SILVA version 132) with a 97% similarity and above required for identification of operational taxonomic units (OTUs). UCHIME (Edgar *et al*. 2011) was also used to identify and remove chimeric fragments. Sequences that could not be directly identified through SILVA, were manually run through NCBI’s BLAST (Madden 2002). We were able to identify all but one OTU with 97% similarity; OTU005 is thus reported as ‘unnamed’ in this study.

### Data analysis

All data, except the 16s rRNA and C:N results, met assumptions of normality and homogeneity of variance. We considered *p*< 0.05 as significant and analyzed data using R version 4.0.3 (R Core Team 2020) and the R packages car (Fox and Weisberg 2019), vegan (Oksanen *et al*. 2020), ggplot2 (Wickham 2016), and entropart (Marcon and Herault 2015). Non metric multidimensional scaling analyses (NMDS) were run using the PAST 3 software (Hammer *et al*. 2001). Due to replication error early in the experiment only the controls and the 300ppm concentration variables were used in statistical difference testing. All GPP and CR data were normalized to mg O_2_ in 120mL per mg of biofilm in 12 hours before being analyzed. C:N data was normalized to mass (mg C/N per mg biofilm). A one way analysis of variance (ANOVA) analysis was used to test for significant differences among treatments and the controls for the GPP, CR, and AFDM. A Kruskal-Wallis test was used to test for significant differences among the C:N ratios. Coefficients of variation were also calculated for individual flumes for each analysis.

Changes in EEA were analysed through both a distance-based redundancy analysis (db-RDA) and three one-way ANOVAs, one per extracellular enzyme. All the data was used within the db-RDA and only the maximum concentrations of each chemical and control were used for analysis in the ANOVA. Where appropriate, we applied a Tukey HSD post-hoc test to the ANOVA output.

Bacterial OTUs were analyzed for alpha and beta diversity as well as differences in the overall bacterial community composition. We used the Chao1 (Chao 1984) and the Shannon-Wiener Diversity Indices to quantify bacterial Alpha diversity (Jost 2007). Chao 1 and Shannon-Wiener results were then analyzed using an ANOVA followed by a Tukey HSD post-hoc test to identify differences in diversity among treatments. Following this, all singletons and doubletons were removed, and the data was rarefied based on the smallest number of sequences identified within a sample. Beta diversity was then analysed using a non-metric multidimensional scaling analysis (NMDS) on all data present to identify differences in the biofilm bacterial community among treatments and controls. An analysis of similarities (ANOSIM) was performed with 999 permutations to identify if these differences were significant. Families and genera were compared graphically, focusing on the top 80% of OTUs identified within the biofilms.

## Results

### Primary productivity and community respiration

Lake control bottles had an average primary productivity of 0.0304 ± 0.0256 mg O_2_ in 120mL and average community respiration of -0.016 ± 0.0177 mg O_2_ in 120mL. We found no significant differences in primary productivity (ANOVA, *F*=0.022, *p*=0.978) and community respiration (ANOVA, *F*=0.148, *p*=0.865) of the treatments versus the controls (Figures 1A, 1B). All treatments showed substantial variation among replicates (Figure 1), with the most pronounced variability being present in respiration. Coefficients of variation were highest for respiration in the controls (1.56, see Table 1) and lowest (0.021) in the rhamnolipid 150 treatment. Rhamnolipid treated systems overall had the lowest variation for CR, while the controls had the highest.

**Table 1.** Coefficients of variation for AFDM, GPP, and CR data. Data collected from samples of biofilm grown in either the surfactant SDS (sodium dodecyl sulphate) or RL (rhamnolipid) for 32 days.

| **Treatment** | **Ash free dry mass** | **Gross primary productivity** | **CR** | **C:N** |
| --- | --- | --- | --- | --- |
| SDS 150 | 0.70 | 0.67 | 0.92 | 0.373 |
| SDS 300 | 0.11 | 0.626 | 1.09 | 0.643 |
| RL 150 | 0.60 | 0.34 | 0.021 | 0.0875 |
| RL 300 | 0.32 | 0.52 | 0.59 | 0.0698 |
| Control | 0.59 | 0.61 | 1.56 | 0.569 |

**Fig. 1.**
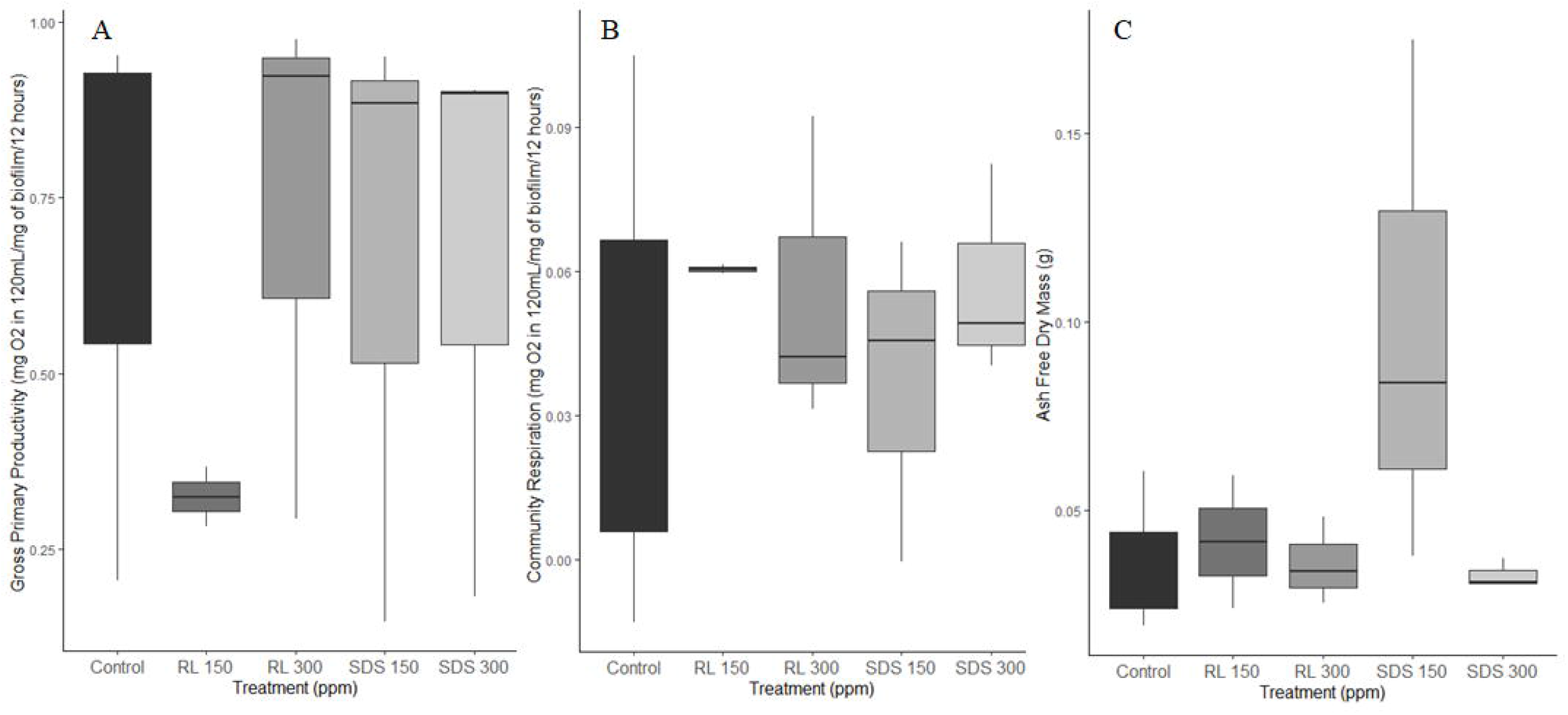
GPP (A), CR (B) and AFDM (C) of microbial biofilms exposed to SDS (sodium dodecyl sulphate) and RL (rhamnolipid) for 32 days at 150 and 300ppm.

### Ash free dry mass and C:N ratios

We found no significant difference among treatments and controls (ANOVA *F*=0.045, *p*=0.957) for AFDM. The coefficients of variation, however, showed differences in variation among the treatments (Table 1), with higher surfactant concentrations decreasing variation (e.g., RL 150 has greater variation than RL 300) (Figure 1C). Control variation was comparable to both low dose surfactant variations. However, C (ANOVA *F*=18.36, *p*=0.0028) and N (ANOVA *F*=5.471, *p*=0.044) content was significantly different by mass among treatments (Figure 2A and Figure 2B). Differences lie between the RL 300 and the control for both C (Tukey HSD *p*=0.0063) and N (Tukey HSD *p*=0.041) content, and between SDS 300 and the control for C (Tukey HSD *p*=0.0036) content.

**Fig. 2.**
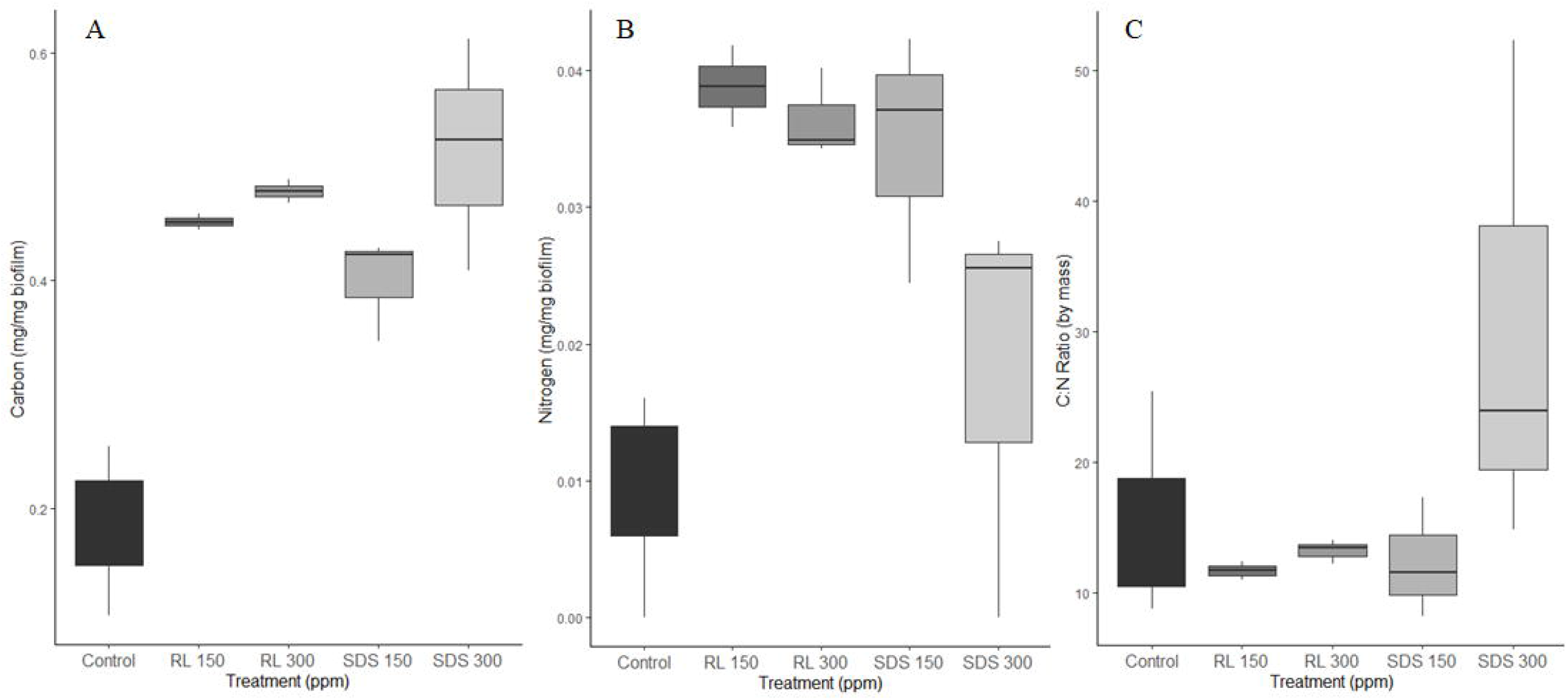
Carbon (A), Nitrogen (B) and C:N ratios (C) of microbial biofilms exposed to SDS (sodium dodecyl sulphate) and RL (rhamnolipid) for 32 days at 150 and 300ppm.

There were also no significant differences among treatments and controls for C:N ratios by mass (Kruskal-Wallis *p*=0.193) present in the biofilms (Figure 2C). There were differences in variation among treatments and controls (Table 1). Rhamnolipid treated flumes had lower variation than either SDS treated flumes or the controls, however within each treatment a higher surfactant concentration increased variation.

### Bacterial community composition

Shannon-Wiener Diversity was significantly different among treatments (ANOVA *F*=71.702, *p*<0.0001, Tukey HSD rhamnolipid and control; *p*=0.043, SDS and control; *p*<0.0001, SDS and rhamnolipid; *p*<0.001) with a higher diversity in the control flumes, second highest in the rhamnolipid flumes, and lowest in the SDS flumes. Significance was driven by surfactant presence and type, but not concentration. The control had the greatest bacterial diversity, while both rhamnolipid treatments had the next highest, and the lowest belonged to both SDS treatments (Figure 3A). Chao1 had no significant differences (ANOVA *F*=3.7748, *p*=0.087).

**Fig. 3.**
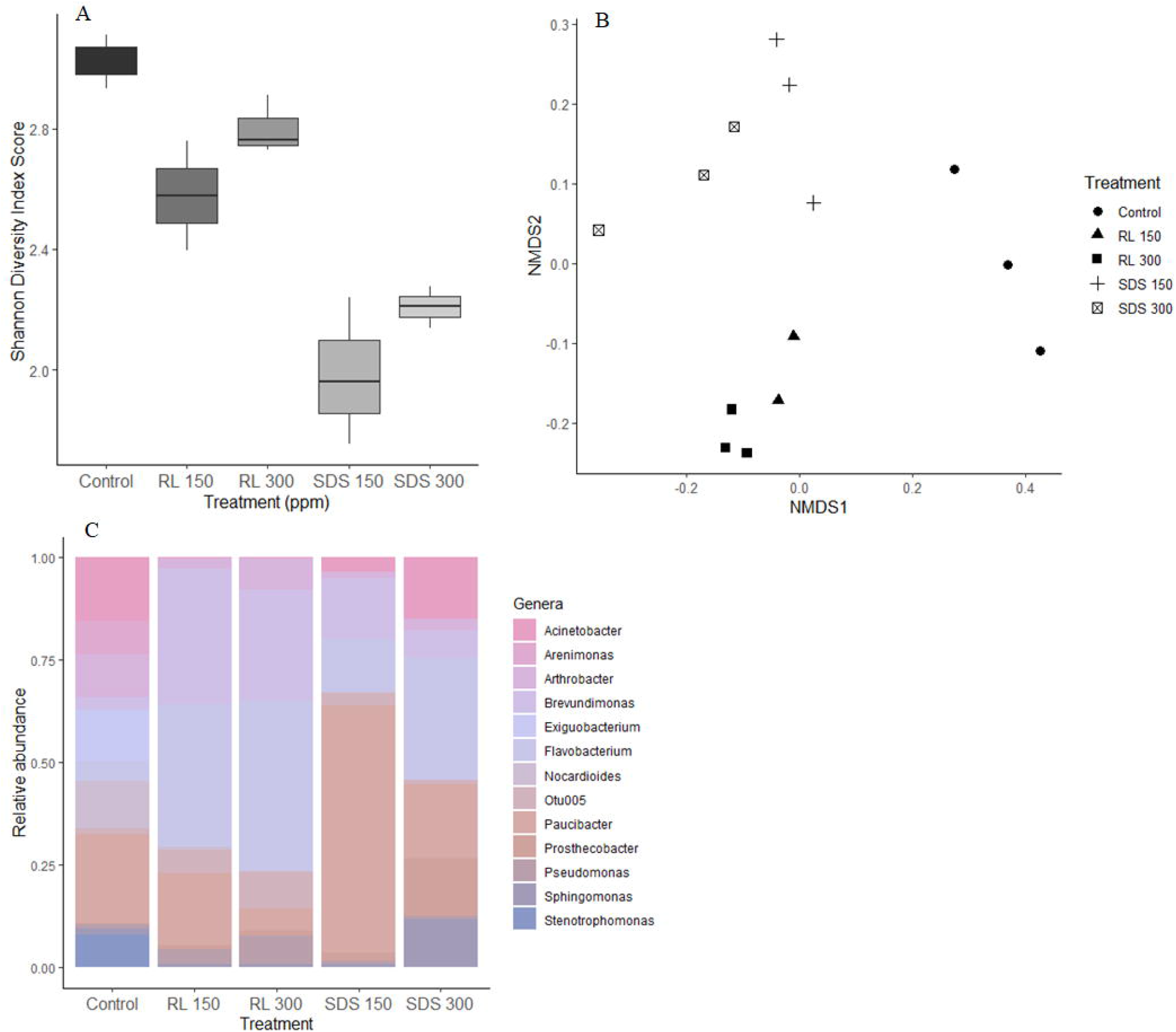
Biofilm bacterial diversity from 16s rRNA sequencing of biofilm exposed to SDS (sodium dodecyl sulphate) or RL (rhamnolipid) after 32 days at 150 and 300ppm. A) shows Shannon Diversity, B) shows an NMDS depicting the community differences in treatments when examining all OTUs present within each sample after rarefication and C) depicts relative abundance of OTUs.

By testing beta diversity and community differences on the rarefied data, we identified distinct groupings between all treatments and controls, as evident by the NMDS (Figure 3B). The ANOSIM results confirmed the differences among both surfactants and the controls (*R*=0.9112, *p*=0.001). Community differences were further identified by looking at only the taxa in the top 80% of all taxa present across all treatments, which was 18 taxa. We identified several genera that were heavily dominant in rhamnolipid-treated flumes. These were: *Brevundimonas spp*., OTU005, *Pseudomonas*, and *Arthrobacter*. In SDS flumes we observed *Brevundimonas terrae, Flavobacterium spp. A*, and in the control flumes *Exiguobacterium, Arenimonas*, and *Stenotrophomona*. We also identified 32 genera only present in the controls, 5 there were exclusive to RL 150, 6 in RL 300, 19 in SDS 150, and 7 in SDS 300 (Figure 3C).

Total OTUs within each treatment and OTUs within the top 80% of all taxa per treatment were also calculated (Table 2). The control flumes had the greatest number of OTUs present, and the greatest number of OTUs within their top 80% of taxa. The SDS and rhamnolipid flumes had similar numbers of total OTUs present, but the SDS flumes had 80% of their total OTUs within half the number of taxa as the rhamnolipids. This indicated a large difference in evenness between treatments, while the total number of OTUs was fairly similar.

**Table 2.** Biofilm OTUs per surfactant treatment and their dominant genera. Biofilms were grown in either SDS (sodium dodecyl sulphate) or rhamnolipid for 32 days.

| Surfactant | Conc. (ppm) | Total OTUs present | OTUs in top 80% per treatment | OTUs lost compared to control | OTUs gained compared to control | Dominant genera in top 80% of all taxa present |  |  |  |  |  |  |
| --- | --- | --- | --- | --- | --- | --- | --- | --- | --- | --- | --- | --- |
| Rhamnolipid | 150 | 145 | 10 | 58 | 25 | *Brevundimonas* | *Otu005* | *Pseudomonas* | *Arthrobacter* | *Flavobacterium B* | *Prosthecobacter* | *Nocardioides* |
| Rhamnolipid | 300 | 159 | 14 | 44 | 36 | *Brevundimonas* | *Otu005* | *Pseudomonas* | *Arthrobacter* | *Flavobacterium B* | *Prosthecobacter* |  |
| SDS | 150 | 176 | 7 | 27 | 37 | *Brevundimonas terrae* | *Acinetobacter* | *Flavobacterium A* | *Paucibacter* | *Flavobacterium B* | *Prosthecobacter* |  |
| SDS | 300 | 130 | 6 | 73 | 23 | *Brevundimonas terrae* | *Acinetobacter* | *Flavobacterium A* | *Prosthecobacter* | *Flavobacterium B* | *Prosthecobacter* | *Sphingomonas* |
| Control | 0 | 203 | 15 | -- | -- | *Exiguobacterium* | *Acinetobacter* | *Arenimonas* | *Stenotrophomonas* | *Nocardioides* |  |  |

### Extracellular Enzyme Activity

According to the db-RDA, EEA differed among all treatments and controls, with the exception of the rhamnolipid treatments. The differences in db-RDA-distances were mainly caused by the differences in EEA of leucine aminopeptidase and phosphatase, although the results were not significant (ANOSIM *F*=2.9907, *p*=0.058). While phosphatase did not have any significant differences among treatments (ANOVA *F*=2.934 *p*=0.129), with the exception of SDS 150 addition of any surfactant decreased the production of phosphatase when compared to the control (Figure 4A). Rhamnolipid additions yielded a significantly higher production of glucosidase, while SDS additions reduced production (ANOVA F=5.741 *p*=0.0404, Tukey HSD *p*=0.034) (Figure 4B). The effects of RL 300 and SDS 300 on leucine aminopeptidase were also significantly different (ANOVA *F*=5.914 *p*=0.038, Tukey HSD *p*=0.032). The addition of rhamnolipid yielded a higher amount of the extracellular enzymes when compared to the control and the addition of SDS reduced production (Figure 4C).

**Fig. 4.**
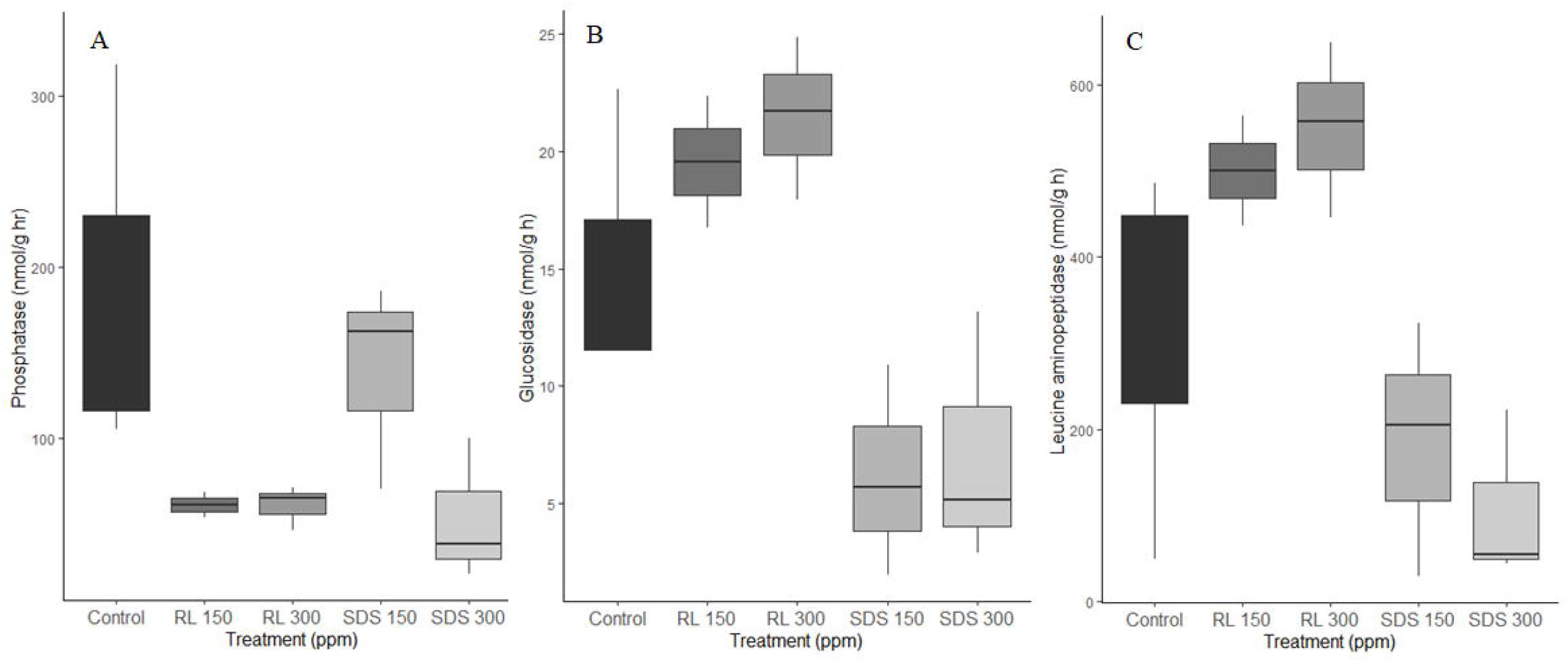
EEA of phosphatase (A), glucosidase (B), and leucine aminopeptidase (C) of biofilms after exposure to either SDS (sodium dodecyl sulphate) or RL (rhamnolipid) for 32 days at 150 and 300ppm. (*indicates significant differences)

## Discussion

Our experiment showed that exposure to biological and synthetic surfactants alters the metabolic activity and the community composition of a mixed-species freshwater biofilm. The effects of SDS were more pronounced than the effects of rhamnolipid. Our experiment is the first study to our knowledge that examines the effects of these two surfactants on a multi-species near-natural freshwater biofilm, which contrasts with the commonly performed single-species studies. This study was also novel as it directly compared the effects of a synthetic and a biological surfactant and explicitly studied for the first time the C:N ratios and C and N contents of biofilms that are exposed to surfactants. These results differ from previous research examining the effectiveness of biosurfactants on biofilm removal in medical (Ceresa et al. 2019; Ceresa et al. 2021b) and industrial (Campos et al. 2013) environments.

### Primary Productivity, Respiration, Ash Free Dry Mass, and C:N ratios

Although our results showed biofilms had no differences in GPP or CR in any treatments or controls, chemical exposure decreased biofilm respiratory activity in previous experiments, including surfactant exposure (Simoes *et al*. 2008) and pharmaceutical exposure (Rosi-Marshall *et al*. 2013). Biofilm volume has also been reduced after a single surfactant exposure event, with decreases of 26.5-98.1% occurring between 22-48 hours after exposure (Mireles II *et al*. 2001; Quinn *et al*. 2013). Typically, biosurfactants, which are amphiphilic polymers, can affect biofilm surface attachment through altering surface wettability (Rodrigues *et al*. 2006). Biosurfactants such as rhamnolipids are able to make cell surfaces more hydrophobic through loss of fatty acids such as lipopolysaccharides (Al-Tahhan *et al*. 2000). Amphiphilic anionic surfactants work similarly through altering surface characteristics and changing solubility of certain compounds (Cserháti *et al*. 2002). Surface changes then can alter a biofilm’s ability to adhere to a surface, reducing growth and development. However, previous work on surfactant effects have also shown that biofilms demonstrate resistance and can recover full respiratory activity 12 hours after the initial exposure (Simoes *et al*. 2008) and biomass 5 hours after initial exposure (Nguyen *et al*. 2020). As our experiment lasted 32 days, we believe that GPP, CR, and AFDM could have stabilized and recovered full respiratory activity as seen in previous studies (Simoes *et al*. 2008; Nguyen *et al*. 2020).

Altered C:N ratios are indicative of changes in microbial productivity (where reduced C:N ratios may yield greater productivity) (Li *et al*. 2016) and biofilm formation (where lower C:N ratios increase biofilm volume) (Thompson *et al*. 2006). As we did not have any alterations in C:N ratios, and no changes in productivity (GPP and CR) or growth (AFDM) among any treatments or controls, our results support these studies. The addition of surfactants provided an added source of C, however as there were consistent C:N ratios across all flumes it can be assumed that our system was C limited. It is also likely that a thickened EPS matrix, which may be used as a defense mechanism in biofilms in response to surfactant exposure, could be another source of extra carbon (Jefferson 2004; Zakaria and Dhar 2020). In contrast, in systems where C is not limiting, an increase in C inputs through surfactants may increase C:N ratios, and therefore negatively affect biofilm productivity and growth.

### Diversity and bacterial community composition

Our controls had taxa present that were minimally identified within the surfactant flumes. These taxa included species (spp.) of *Exiguobacterium, Acinetobacter, Arenimonas*, and *Stenotrophomonas. Exiguobacterium* is currently researched as an emerging human pathogen and while susceptible to antibiotics appears to easily develop antibiotic resistance (Chen *et al*. 2017). *Acinetobacter* contains species such as *A. baumanni* that are very well known as human pathogens and that demonstrate high antibiotic resistance (Rice 2008; Tewari *et al*. 2018). However, it appears that some species may not have resistance to biological surfactants as its presence in rhamnolipid treated flumes was extremely low, although certain species of *Acinetobacter* had an increase in abundance within the SDS treated flumes. Similar results appear with *Stenotrophomonas* as it has human pathogenic potential, and species that have developed multi drug resistance, but appears to be sensitive to both biological and synthetic surfactants (Chang *et al*. 2015). In our study the addition of either surfactant created an environment unsuitable for these bacteria, demonstrating how surfactant exposure can alter biofilm community composition and potentially help to control certain pathogenic species that are demonstrating resistance to antibiotics. The control flumes had the highest amount of OTUs present (*n*=203) and the highest diversity, which indicates that both the diversity and abundance of bacteria present in the biofilms decreases after surfactant exposure.

Rhamnolipids are studied for their ability to reduce pre-existing biofilms and prevent future biofilm growth with some success (Quinn *et al*. 2013; Dusane *et al*. 2010; De Rienzo *et al*. 2015; Satpute *et al*. 2018). Current research has shown that rhamnolipid exposure on soil bacteria can shift community composition, although many studies do not examine differences to the genus level (Lu *et al*. 2019; Wei *et al*. 2020; Akbari *et al*. 2021). Research has also shown that aquatic microbial communities will shift in response to an outside contaminant with an increased abundance of bacteria that are not sensitive to that contaminant or can use it as a C source (Hay *et al*. 2001; Sumpono *et al*. 2003; Seghers *et al*. 2003; Drury *et al*. 2013). Our study showed that, much like soil communities and aquatic communities exposed to other contaminants, aquatic biofilms experience significant community changes after exposure to rhamnolipids.

Further examination into the bacterial genera indicate large increases in a *Brevundimonas* spp., *Pseudomonas* spp., *Arthrobacter* spp., and OTU005 when compared to all other treatments and to the controls. One OTU of *Arthrobacter* spp. was only present within the rhamnolipid flumes, with the exception of one control flume that had low levels of that particular OTU. *Brevundimonas* spp., who are opportunistic pathogens (Ryan and Pembroke 2018) have experienced accelerated growth in the presence of rhamnolipids before when testing the effects of rhamnolipids on increasing bioremediation in soils (Lu *et al*. 2019). It is likely that these genera were able to use rhamnolipid as a C source due to the large C quantities within rhamnolipids (Mnif *et al*. 2018). Their adaptability may explain why they were able to increase in abundance over time when compared to the other genera present.

SDS caused the largest reduction in microbial diversity when compared with the other treatments and controls. SDS has been researched and used for its ability to degrade biofilms in applications where biofilms are undesirable (Izano *et al*. 2007; Ueda *et al*. 2019; Nguyen *et al*. 2020). This is particularly due to their ability to bind to hydrophobic protein components, introducing a negative charge to the structure which unfolds the bound protein (Gudiksen *et al*. 2006; Fong and Yildiz 2015). While rhamnolipids are also able to denature proteins, they are considered weak binders, and typically have a reduced denaturing ability when compared to many other surfactants (Otzen 2017). This may explain why biofilms exposed to SDS had lower diversity than those in the rhamnolipid treatments or the controls. SDS also had the lowest amount of OTUs within the top 80% of taxa present in the treatments. We believe that the low diversity, and community domination by few taxa, demonstrates that SDS exposure creates an inhospitable environment to many potential specialist taxa. The few remaining and insensitive taxa can then thrive as more resources are available. Following our findings, these taxa would include *Brevundimonas terrae*, and an *Acinetobacter* and *Flavobacterium spp*. While the rhamnolipid treated flumes also had less diversity than the controls, the rhamnolipid treatments still appeared to favor more taxa and a higher diversity than the SDS treated flumes.

### Extracellular Enzyme Activity

Our experiment showed that substantial changes in biofilm EEA can occur after surfactant exposure. Although phosphatase activity, or the separation of phosphate groups from a larger compound such as an acid or protein (Margalef *et al*. 2017), generally decreased after surfactant exposure, we did have similar levels of activity between the control and SDS 150 flumes. Examination of the bacterial communities within SDS 150 flumes and control flumes show several parallel genera present that are not as common in the other systems, including *Nocardioides*. Treatments were laid out randomly across our experimental flumes within the experimental setup, and water was allocated to all systems from the same source, which minimized potential experimental alterations by relative location. We thus assume that all changes in community composition and EEA refer to the actual biofilm response to treatments. The presence of a low concentration SDS allowed for the development of bacterial genera like *Nocardioides* which typically have both acid and alkaline phosphatase (Xie *et al*. 2017). However, the addition of high concentration SDS, or rhamnolipid at either a high or low concentration, reduced phosphatase activity.

Beta-glucosidase (rate-limiting enzyme that converts cellulase to glucose (Singh *et al*. 2016)) and leucine aminopeptidase (enzyme that aids in the hydrolysis of leucine and other N-terminal residues at the end of peptides and proteins (Matsui *et al*. 2006)) activities were also affected by surfactant exposure. Previous research has shown that rhamnolipids can act as an enhancer for the enzyme beta-glucosidase (Zhang *et al*. 2009; Yao *et al*. 2020). Rhamnolipids are able to protect proteins and limit the degradation of cellulase (Zhang *et al*. 2009; Wang *et al*. 2011). Yin *et al*. (2019) showed that the presence of rhamnolipid can increase genes related to leucine and amino acid synthesis. The addition of rhamnolipid to our flumes likely protected cellulase and leucine in the system which led to enhanced beta-glucosidase and leucine aminopeptidase activity. Beta-glucosidase is also known for its ability to degrade *Pseudomonas aeruginosa*, a rhamnolipid producing bacteria (Banar *et al*. 2019; Pirlar *et al*. 2020). The flumes treated with rhamnolipid had a higher abundance of *P. aeruginosa* when compared with the other flumes.

We assume therefore, that the higher beta-glucosidase levels within the rhamnolipid flumes were likely caused by a combination of the presence of rhamnolipid to act as an enhancer, and more abundant *P. aeruginosa*. However, we observed somewhat contrasting results in the SDS treated flumes, where a decrease in enzyme activity occurred. SDS is an anionic surfactant that can be used in denaturation of polypeptides (Bhuyan 2009; Schlager *et al*. 2012). SDS is commonly used in an analysis called SDS-polyacrylamide gel electrophoresis (SDS-PAGE) where an SDS infused gel is used to separate proteins and enzymes based on their molecular weight. This analytical method has been used with experiments examining beta-glucosidase previously (Laemmli 1970; Bai *et al*. 2013), which demonstrates that SDS is able to denature beta-glucosidase. It is possible that the denaturing ability of SDS has degraded beta-glucosidase and leucine aminopeptidase, leading to reduced enzyme activity, and likely also reduced biofilm metabolic activity.

Our experiment showed that both the biologic surfactant rhamnolipid and the synthetic surfactant, SDS, can significantly affect a mixed freshwater microbial biofilm in terms of community composition and EEA. Our original hypotheses predicted changes to bacterial biodiversity, and biofilm metabolic activity after exposure to either surfactant. Rhamnolipids acted as an enhancer for beta-glucosidase and leucine aminopeptidase, increasing EEA within the biofilm, while SDS likely denatured the enzymes and reduced EEA within the biofilm. While there were no differences in GPP and CR, EEA changes can be used as a proxy for community-scale metabolic activity within the biofilms. There were alterations in biofilm metabolic activity post surfactant exposure, supporting one of our original hypotheses. Both surfactants also decreased the diversity of the biofilm bacterial community, which supports our other original hypothesis. There were unique genera in the biofilms of each treatment, including the controls which demonstrates that the presence of either surfactant creates an environment that is inhospitable to sensitive taxa. Although we saw differences in EEA and diversity, there were no differences in GPP and CR, potentially demonstrating functional redundancy within mixed microbial biofilms. These results highlight several important areas of concern. Firstly, rhamnolipids may not be a suitable alternative to SDS for use environmental biofilm control because of its lower effectiveness in open environmental settings. Secondly, in natural ecosystems where biofilms are necessary for key ecosystem processes, such as degradation of organic matter, we have shown that the presence of these surfactants, (for example through the release from WWTPs) may lead to negative effects on microbial diversity and metabolic activity within biofilms. Finally, our results demonstrate that functional redundancy may exist within diverse biofilms. Surfactant exposure altered community composition and EEA, but the overall biofilm did not experience any negative respiratory effects. Therefore, while the function of the biofilm may be altered, enough bacteria are still taking part in respiratory cycles that the overall metabolic activity of the biofilm is not disrupted.

## Acknowledgements

We thank Gertraud Steniczka and Hermann Hofreiter for help in the laboratory and with setting up the experiment.

## Statements and Declarations

### Ethics approval and consent to participate

Not applicable

### Consent for publication

Not applicable

### Availability of data and materials

Data can be accessed at https://zenodo.org/record/5153485#.YQgBzEAo9PY. Access is restricted and must be requested from the page to prevent work from being copied before publishing. After publishing work will be publicly accessible.

### Author’s contributions

SG, WH, LC, IB, and JS were all equally involved in the design, execution, and data analysis within the study. SG wrote the first draft of the manuscript with the exception of the extracellular enzyme analysis methodology which was written by LC. All authors contributed to commenting on and editing the manuscript and have read and approved of the final manuscript.

